# Longitudinal mapping of cortical change during early adolescence associated with prenatal tobacco and/or alcohol exposure in the Adolescent Brain Cognitive Development^SM^ Study

**DOI:** 10.1101/2024.08.29.610335

**Authors:** Andrew T. Marshall, Shana Adise, Eric C. Kan, Elizabeth R. Sowell

**Affiliations:** Children’s Hospital Los Angeles, Los Angeles, California, 90027, United States of America; University of Southern California, Los Angeles, California, 90027, United States of America

## Abstract

**Importance:** The effects of prenatal alcohol (PAE) and tobacco exposure (PTE) on adolescent neuroanatomical development are typically evaluated cross-sectionally. It is unclear if observed effects persist throughout life or reflect different developmental trajectories.

**Objective:** To determine how PAE and PTE are associated with cortical structure and development across two timepoints in early adolescence.

**Design:** Observational, longitudinal analyses of data within the Adolescent Brain Cognitive Development□ Study

**Setting:** 21 study sites in the United States

**Participants:** 5,417 youth participants, aged ∼9-12 years old

**Exposures:** PAE and PTE based on caregiver (self) reports of alcohol/tobacco use during pregnancy, before and after pregnancy recognition.

**Main Outcomes and Measures:** Cortical thickness (mm) and cortical surface area (mm^2^) measured approximately 2 years apart in early adolescence, across 68 bilateral cortical regions.

**Results:** At baseline data collection, youth participants were ∼9.9 years old (*SD*=0.6). At the second neuroimaging appointment, youth participants were ∼11.9 years old (SD=0.6). When modelling cortical thickness, we controlled for individuals’ whole-brain volume; when modelling cortical surface area, individuals’ total surface area. Cortical thickness generally declined with age. Cortical surface area either expanded or contracted with age, depending on region. PAE had minimal effects on cortical structure (main effects) and development (PAE×Age interactions). PTE had robust effects on cortical thickness and was associated with faster rates of cortical thinning in several regions within the frontal lobe. Post hoc analyses on (1) the effects of PTE for those who continued tobacco use after pregnancy recognition and (2) the effects of PTE in those who did not also use alcohol revealed weaker effects.

**Conclusions and Relevance:** PTE had robust effects on neuroanatomical structure and longitudinal development, particularly cortical thickness. Analyzing developmental cortical trajectories informs how PTE and/or PAE not only affects cortical structure but how it develops long after those prenatal exposures occurred. Future analyses involving cotinine biomarkers of PTE would enhance the temporal resolution of the ABCD Study^**®**^’s PTE-related queries of tobacco use before and after learning of the pregnancy.

**Key Points:**

- How does prenatal tobacco and/or alcohol exposure affect brain structure and longitudinal development in early adolescence?
- Prenatal tobacco exposure was robustly associated with faster rates of cortical thinning in the frontal lobe; prenatal alcohol exposure had more minimal effects.
- Prenatal substance exposure, especially prenatal tobacco exposure, not only affects cortical structure but how it develops long after those prenatal exposures occurred.

## Introduction

Environmental factors can affect healthy brain development during critical developmental windows, like early childhood and adolescence.^1^ We showed that the risk of lead exposure more detrimentally impacted brain structure in youth in more socioeconomically disadvantaged families.^2^ Such neurotoxicity is also associated with prenatal alcohol (PAE) and tobacco exposure (PTE).^3^ Despite product labels advising against alcohol/tobacco use while pregnant,^4^ many individuals do so (either before or after knowing of the pregnancy).^5,6^ PAE is associated with smaller total/regional brain volumes and poorer connectivity.^7^ Similarly, PTE is associated with reduced cortical thickness, surface area, and/or volume across various brain regions.^3,8-15^ While research has investigated the cross-sectional associations of PAE and PTE,^3,11^ less is known about neuroanatomical developmental trajectories given PTE and/or PAE long after they occur, which could elucidate, for example, whether individuals with PTE have thinner cortices initially and/or show faster rates of cortical thinning during childhood/adolescent brain development.

To address these unresolved questions, we analyzed data from the Adolescent Brain Cognitive Development□ Study (hereafter, “ABCD”), a 10-year, longitudinal research study occurring at 21 U.S. study sites.^17^ Recent work showed that ABCD participants with PTE had smaller cortical surface areas and volumes at ABCD’s first two neuroimaging timepoints (i.e., early adolescence).^12^ Another ABCD study evaluated whether main effects of PAE and PTE on brain/behavior were consistent between those timepoints,^18^ but neither specifically considered the change over time between assessments. Here, we determined whether PAE and PTE moderated developmental change in cortical thickness and surface area across those timepoints, hypothesizing (given past research) that PAE and PTE would be associated with greater cortical thinning (i.e., reduced and more rapidly decreasing thickness) and faster surface contraction (i.e., reduced and more rapidly decreasing area). Then, post hoc analyses determined whether significant PAE/PTE×Age interactions were present in (1) children of individuals who continued to use alcohol and/or tobacco after having learned of the pregnancy and/or (2) children of individuals who reported alcohol but not tobacco use or tobacco but not alcohol use.

## Methods

### Participants

ABCD enrolled ∼11,900 9- and 10-year-old youth primarily using school-based enrollment.^19^ Our data came from the 2021 ABCD 4.0 data release (doi: 10.15154/1523041; https://data-archive.nimh.nih.gov/abcd), which included baseline data for 11,876 youth and two-year follow-up data for 10,414 youth (i.e., the two neuroimaging timepoints in the 4.0 release). Centralized IRB approval was obtained from the University of California, San Diego. Study sites obtained approval from their local IRBs. Caregivers provided written informed consent and permission; youth provided written assent. Data collection/analyses complied with all ethical regulations.

### Prenatal Exposure, Birth Weight, and Demographics

At baseline, caregivers reported the biological mother’s substance use (1) before knowing of pregnancy but when they could have been pregnant with the youth participant and (2) after knowing of the pregnancy. For both, response options were “Yes”, “No”, and “Don’t know” (DK). We focused on alcohol and/or tobacco use. Caregivers also reported their child’s birthweight. Sex-assigned-at-birth was included in all publicly downloadable data tables.

### Neuroimaging

ABCD neuroimaging collection and processing procedures are thoroughly described.^20^ We analyzed the thickness (mm) and surface area (mm^2^) of 34 bilateral cortical regions [derived using FreeSurfer v.7.1.1 Desikan-Killiany atlas on acquired T_1_w structural magnetic resonance imaging (sMRI) volumes].^21,22^ Whole-brain volume, intracranial volume, mean cortical thickness, and total surface area were also analyzed. Neuroimaging parameters are available: https://abcdstudy.org/images/Protocol_Imaging_Sequences.pdf.

### Statistics

Per ABCD’s neuroimaging data-release notes, 8 participants were excluded, as were 209 participants whose T_1_w data were not recommended for analysis (remaining *n*=11,659). We removed 4,227 participants who did not have baseline and two-year sMRI data (remaining *n*=7,432) and 736 participants who had outlying/missing birthweight data (remaining *n*=6,696). (Birthweight was analyzed because PTE is associated with reduced birthweight^23^ and smaller childhood/adolescent brain volume,^9^ which are also associated with each other.^24^) Because ABCD includes siblings, we controlled for family relatedness by only including singletons or one sibling per family; we prioritized siblings based on PAE/PTE and data completeness (yes/no responses to PTE/PAE before/after learning of the pregnancy: PTE/Before, PTE/After, PAE/Before, PAE/After). If siblings had identical sets of yes/no/DK responses, we randomly selected one sibling from the corresponding family groups using MATLAB’s *datasample* function (seed=1) [MATLAB Version: 9.13.0.2126072 (R2022b) Update 3]. Otherwise, we included the sibling based on (1) more “yes” responses to PTE/After and PAE/After pregnancy recognition, (2) more “yes” responses total, and (3) fewest DK responses (remaining *n*=5,713).

Participants were separately categorized into PAE and PTE groups (Table 1). Non-PAE and Non-PTE designations were for participants whose caregivers reported not using alcohol or tobacco products before or after learning/knowing of the pregnancy; PAE and PTE designations, for participants whose caregivers did report such use. Given a priori plans to conduct post hoc analyses on PAE/PTE after learning/knowing of the pregnancy, we excluded participants whose caregivers were unaware of the pregnant person’s alcohol/tobacco use during that period, regardless of what they reported before learning/knowing of the pregnancy. Thus, 5,417 participants remained in analyses (PAE: *n*=1,500; Non-PAE: *n*=3,917; PTE: *n*=739; Non-PTE: *n*=4,678).

**Table 1.**
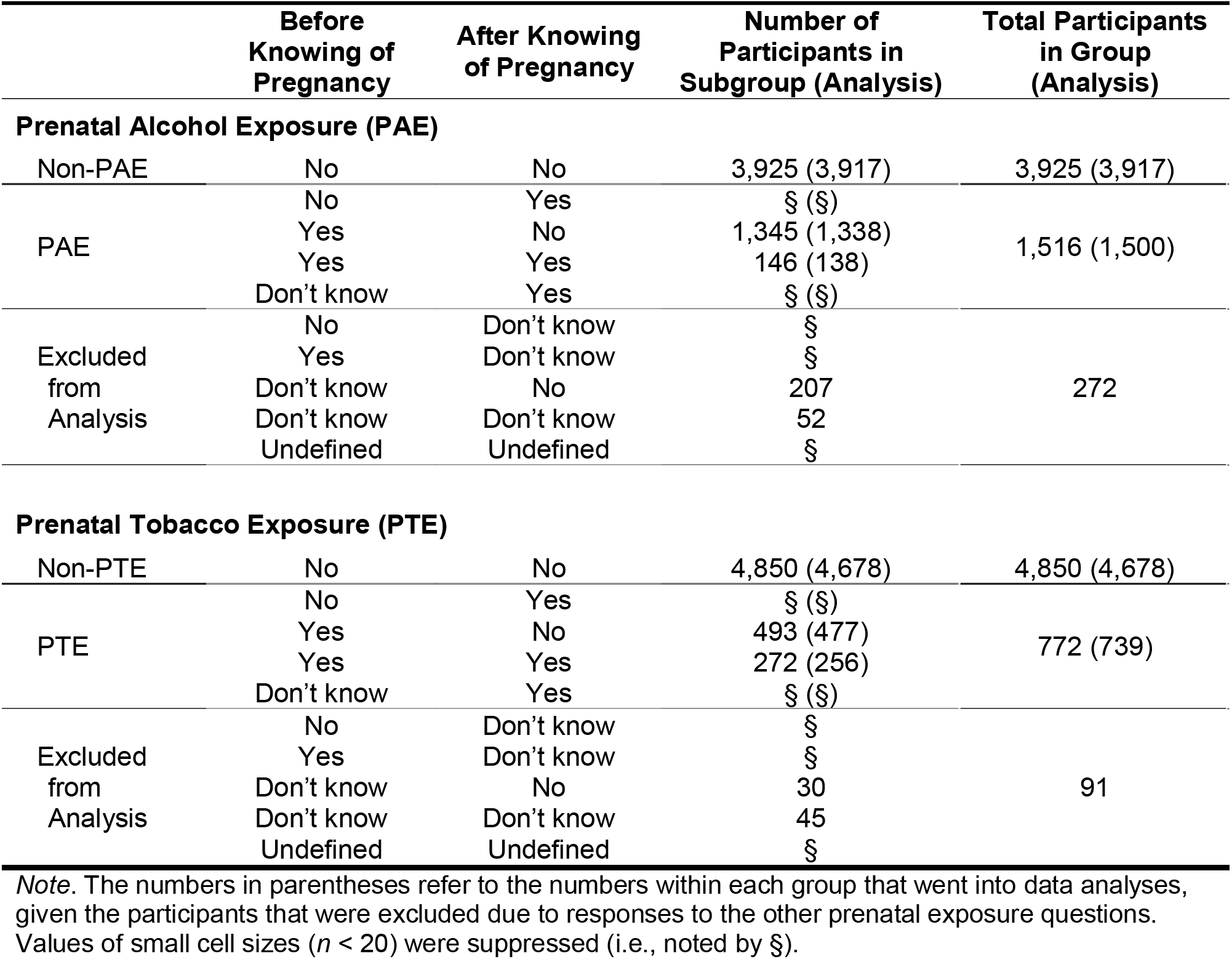
Prenatal alcohol and prenatal tobacco exposure group criteria based on caregivers’ responses.

We first determined a whole-brain covariate for analyzing longitudinal change in regional thickness and surface area.^25^ For both, across 68 bilateral regions, we ran 4 linear mixed-effects models:

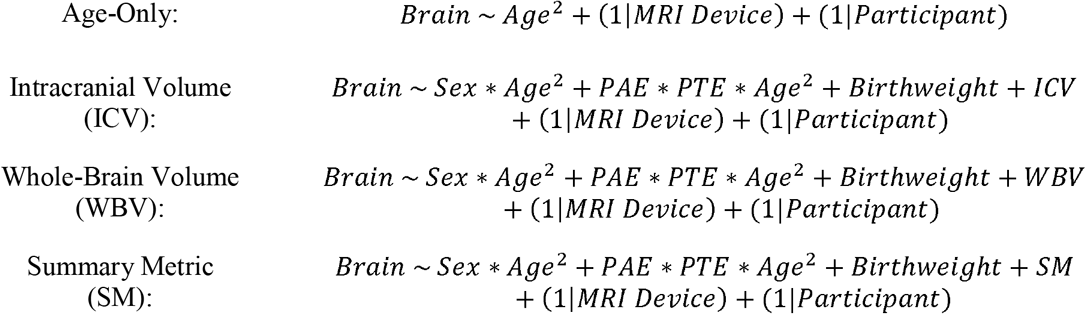

Age (months), birthweight (total ounces), ICV, WBV, and SM (mean thickness or total area) were continuous, mean-centered predictors. While we focused on linear interactions, *Age*^2^ was included in all models given curvilinear changes in brain structure over longer stretches of adolescence.^26^ Sex, PAE, and PTE were effect-coded categorical factors with male/female, Non-PAE/PAE, and Non-PTE/PTE, respectively, coded as -1/+1. By-MRI-device and by-participant intercepts were random effects. All models used a random initial value for iterative optimization. For each region-by-region analysis, participants’ data were considered outliers (and excluded) if either of their datapoints exceeded more than three scaled median absolute deviations away from the median.^27^ This resulted in between 10 and 241 participants excluded from thickness analyses (*n*’s=5,176-5,407); from surface-area analyses, between 4 and 107 participants (*n*’s=5,310-5,413).

Separately, for thickness and surface area, the more optimal whole-brain covariate was selected based on two criteria: (1) a Pearson’s-*r* correlation between the fixed-effects coefficients (across 68 ROIs) from the age-only models and those from the ICV, WBV, and SM models; higher correlations reflected greater similarities between two sets of coefficients; (2) the residual total sum-of-squares (TSS) between each of the ICV, WBV, and SM models and the age-only model (sum of the squared differences between the coefficients from each of the former and the latter); smaller values reflected greater similarities between sets of coefficients. The chosen set of models were used to evaluate PTE/PAE×Age interactions on brain structure.

Effect sizes for age are represented by partial correlation coefficients (*r*_*p*_), which account for all other predictors.^28^ For consistency, the strengths of modelled interactions and categorical factors were calculated similarly. The 95% CIs of effect sizes were derived from the sample variance of the partial correlation.^29^ The Benjamini-Hochberg^30^ false-discovery-rate (FDR) algorithm was used to correct for multiple comparisons. eTables 1-52 show output for associations of and interactions between age, PAE, and PTE from thickness and surface-area models controlling for ICV, WBV, and SM.

## Results

### Sample Characteristics

At baseline, youth participants were ∼9.9 years old (*SD*=0.6), and, at the two-year appointment, ∼11.9 years old (SD=0.6). Our sample was similar to the entire ABCD cohort but, proportionally, was more likely to live in more socioeconomically advantaged families and be reported by their caregivers as White and not Hispanic/Latino/a/x (Table 2).

**Table 2.** Participant Demographics.

|  | ABCD Cohort –<br>Baseline (%) | Sample in This<br>Report - Baseline (%) |
| --- | --- | --- |
| **Youth Sex** |  |  |
| Male | 6,196 (52.2%) | 2,912 (53.8%) |
| Female | 5,680 (47.8%) | 2,505 (46.2%) |
| **Annual Household Income** |  |  |
| <\$50K (Low) | 3,223 (27.1%) | 1,381 (25.5%) |
| \$50-100K (Mid) | 3,071 (25.9%) | 1,509 (27.9%) |
| >\$100K (High) | 4,564 (38.4%) | 2,123 (39.2%) |
| Missing/Undefined | 1,018 (8.6%) | 404 (7.5%) |
| **Caregiver Education** |  |  |
| ≤12th Grade/No Diploma | 593 (5.0%) | 223 (4.1%) |
| High-school Graduate/GED or<br>Equivalent | 1,132 (9.5%) | 456 (8.4%) |
| Some College, No<br>Degree/Associate's Degree | 3,079 (25.9%) | 1,416 (26.1%) |
| Bachelor's Degree | 3,015 (25.4%) | 1,427 (26.3%) |
| Master's/Professional Degree,<br>Doctorate | 4,043 (34.0%) | 1,888 (34.9%) |
| Missing/Undefined | § (§%) | § (§%) |
| **Youth Race** |  |  |
| American Indian / Alaska Native | 62 (0.5%) | 24 (0.4%) |
| Asian | 275 (2.3%) | 95 (1.8%) |
| Black | 1,869 (15.7%) | 724 (13.4%) |
| Native Hawaiian / Pacific<br>Islander | § (§%) | § (§%) |
| Other | 1,959 (16.5%) | 867 (16.0%) |
| White | 7,524 (63.4%) | 3,640 (67.2%) |
| Missing/Undefined | 171 (1.4%) | 60 (1.1%) |
| **Youth Ethnicity** |  |  |
| Hispanic | 2,411 (20.3%) | 1,048 (19.3 %) |
| Not Hispanic | 9,312 (78.4%) | 4,314 (79.6%) |
| Missing/Undefined | 153 (1.3%) | 55 (1.0%) |
| **Total** | **11,876 (100%)** | **5,417 (100%)** |

*Note.* The data in the “ABCD Cohort” column refer to those derived from the 4.0 ABCD Study Data Release (October 2021). The “Other” Race/Ethnicity category includes those who identified “Other Race” and those who identified more than one race (i.e., multiracial). Values of small cell sizes ( $n < 20$ ) were suppressed (i.e., noted by §).

### Prenatal Exposure

#### Whole-brain covariates

The linear coefficients for age from age-only cortical-thickness models were more comparable to those from the WBV and ICV models [WBV: *r*>.99, *p*<.001, TSS=1.14×10^−6^; ICV: *r*>.99, *p*<.001, TSS=1.25×10^−6^] than models controlling for mean thickness (*r*=.89, *p*<.001, TSS=1.11×10^−4^]. Accordingly, we controlled for WBV in cortical-thickness models. In surface-area models, we controlled for total area (SM: *r*>.99, *p*<.001, TSS=0.72; WBV: *r*=.96, *p*<.001, TSS=11.35; ICV: *r*=.72, *p*<.001, TSS=145.92).

#### Cortical development: Main effect of age

There were FDR-corrected age-associated decreases in cortical thickness in 65 regions (Figure 1A). For surface area, 56 regions showed FDR-corrected age-associated changes, 34 of which expanded (i.e., posterior-frontal, superior-temporal lobes), with 22 contracting (i.e., parietal, occipital, inferior-temporal lobes) (Figure 1B).

**Figure 1.**
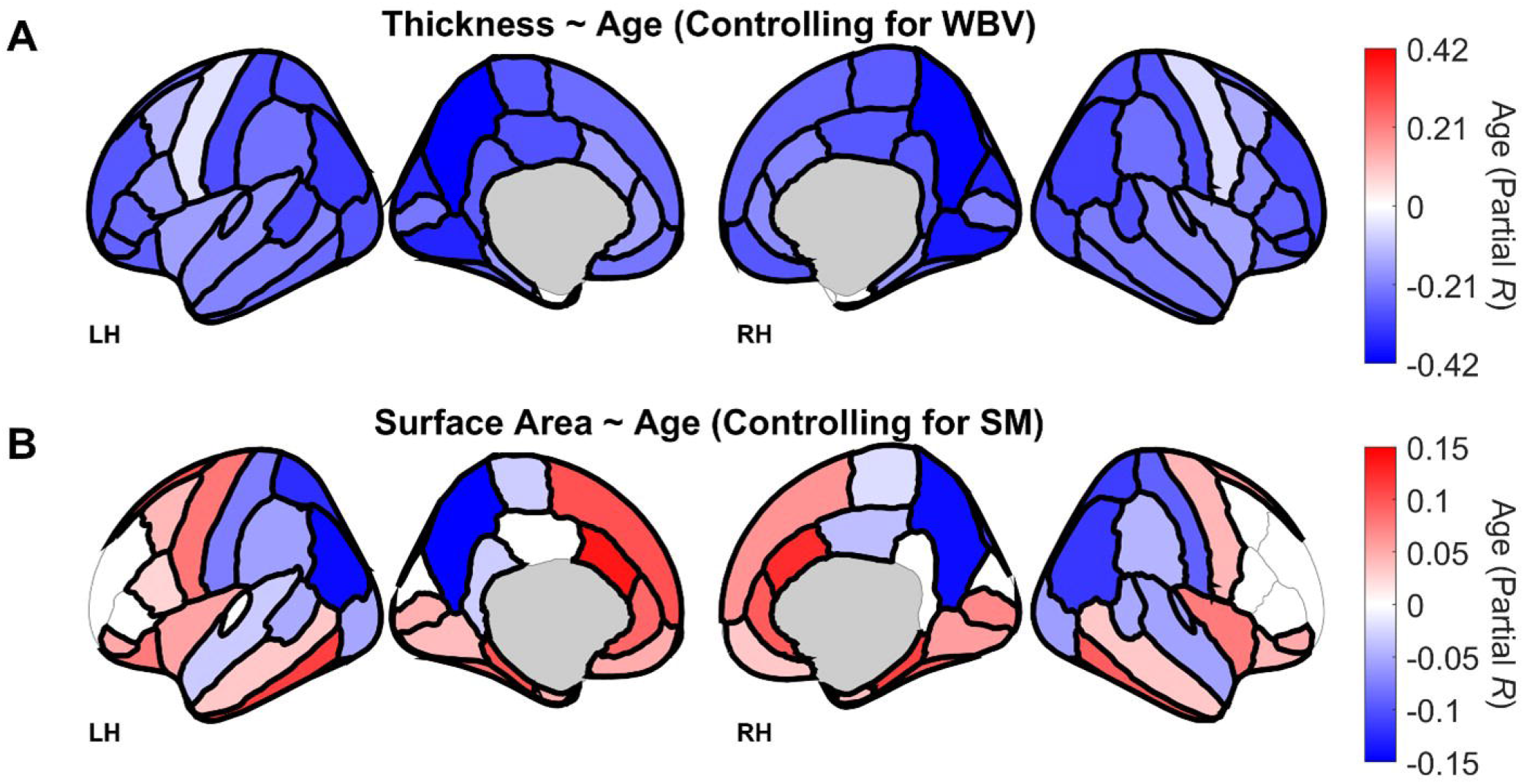
Associations between participant age and brain structure. Regions are color coded with respect to the color bars on the immediate right of each panel. (Note the different scales of the color bars.) Red- or blue-shaded regions indicate that those associations had *p*-values less than .05; regions that have thick borders indicate that those associations also passed false discovery rate (FDR) correction. These images were generated in MATLAB using data from the *ggseg* toolbox in R.^42^ LH = left hemisphere. RH = right hemisphere. SM = total surface area. WBV = whole-brain volume.

#### PAE: Main effects and interactions with age

PAE’s effects on thickness were minimal, with only the right-parahippocampal cortex being thicker (main effect), *p*=.017 (eTable 16), and the left-caudal-anterior-cingulate thinning more slowly in those with versus without PAE, *p*=.007 (eTable 18) (PAE×Age); neither passed FDR correction. Those with PAE (main effect) showed greater surface areas in 3 regions (left-inferior-temporal, right-middle-temporal, left-fusiform), *p*s≤.017, and smaller areas in 4 regions (right-postcentral, left-inferior-parietal, left-frontal-pole, left-medial-orbitofrontal), *p*s≤.050; none passed FDR correction (eTable 23).

There were minimal PAE×Age interactions on surface area (left-entorhinal, left-interior-temporal, left-lateral-orbitofrontal, right-pericalcarine, *p*s≤.031), none passing FDR correction (eTable 25).

#### PTE: Main effects

Those with PTE had thinner cortices in 8 regions (bilateral-parahippocampal, bilateral-precentral, bilateral-paracentral, left-lateral-orbitofrontal, left-temporal-pole), *p*s≤.043; the bilateral-parahippocampal and left-lateral-orbitofrontal cortical effects passed FDR correction (eTable 17). Those with PTE showed thicker cortices in 5 regions (bilateral-pars-opercularis, right-transverse-temporal, right-pars-triangularis, right-caudal-anterior cingulate), none passing FDR correction, *p*s≤.045 (eTable 17) (Figure 2A). PTE was associated with reduced surface area in 6 regions (bilateral-precentral, right-posterior-cingulate, left-pericalcarine, right-entorhinal, left-postcentral) and increased area in 5 regions (bilateral-pars-triangularis, bilateral-insula, left-superior-frontal), *p*s≤.040, which did not pass FDR correction (eTable 24) (Figure 2B).

**Figure 2.**
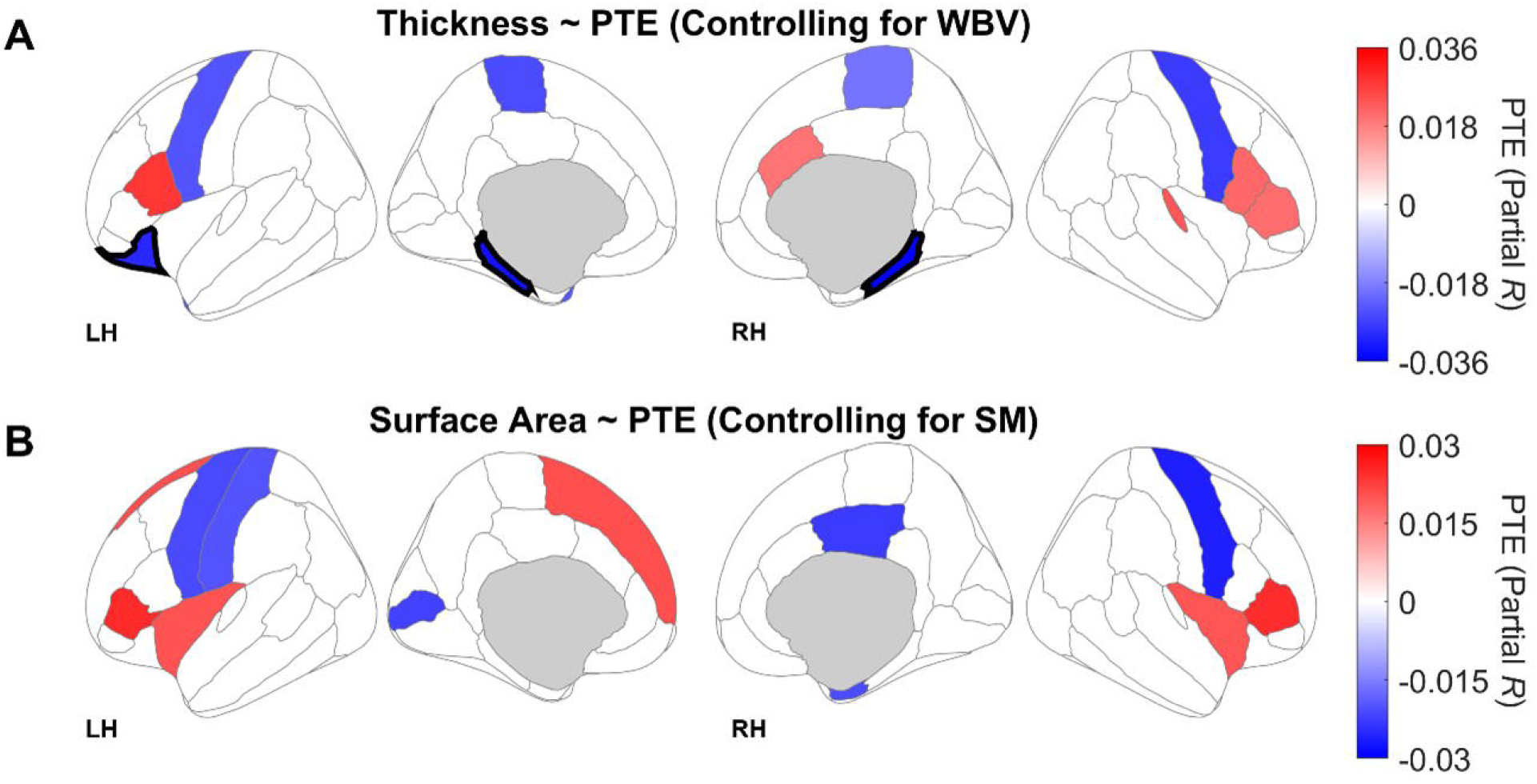
Associations between prenatal tobacco exposure (PTE) and brain structure. Regions are color coded with respect to the color bars on the immediate right of each panel. (Note the different scales of the color bars.) Red- or blue-shaded regions indicate that those associations had *p*-values less than .05; regions that have thick borders indicate that those associations also passed FDR correction. Blue-shaded regions reflect thinner cortices (A) or smaller surface areas (B) in those with PTE. Red-shaded regions reflect thicker cortices (A) or larger surface areas (B) in those with PTE. These images were generated in MATLAB using data from the *ggseg* toolbox in R.^42^ LH = left hemisphere. RH = right hemisphere. SM = total surface area. WBV = whole-brain volume.

#### PTE: Interactions with age

There was a robust pattern of PTE being associated with steeper rates of cortical thinning, with there being FDR-corrected PTE_×_Age interactions in 11 frontal regions (bilateral-rostral-middle-frontal, bilateral-superior-frontal, bilateral-medial-orbitofrontal, bilateral-rostral-anterior-cingulate, right-pars-orbitalis, left-frontal-pole, right-pars-triangularis) and 2 temporal regions (right-banks-of-the-superior-temporal-sulcus, left-inferior-temporal), *p*s≤.005; 9 additional regions (including 3 frontal and 4 temporal regions) showed non-FDR-corrected PTE_×_Age interactions, *p*s≤.043 (eTable 19) (Figure 3). There were PTE_×_Age interactions on surface area in 5 regions (left-supramarginal, right-lateral-orbitofrontal, left-frontal-pole, left-paracentral, left-pericalcarine), *p*s≤.038, but they did not pass FDR correction (eTable 26).

**Figure 3.**
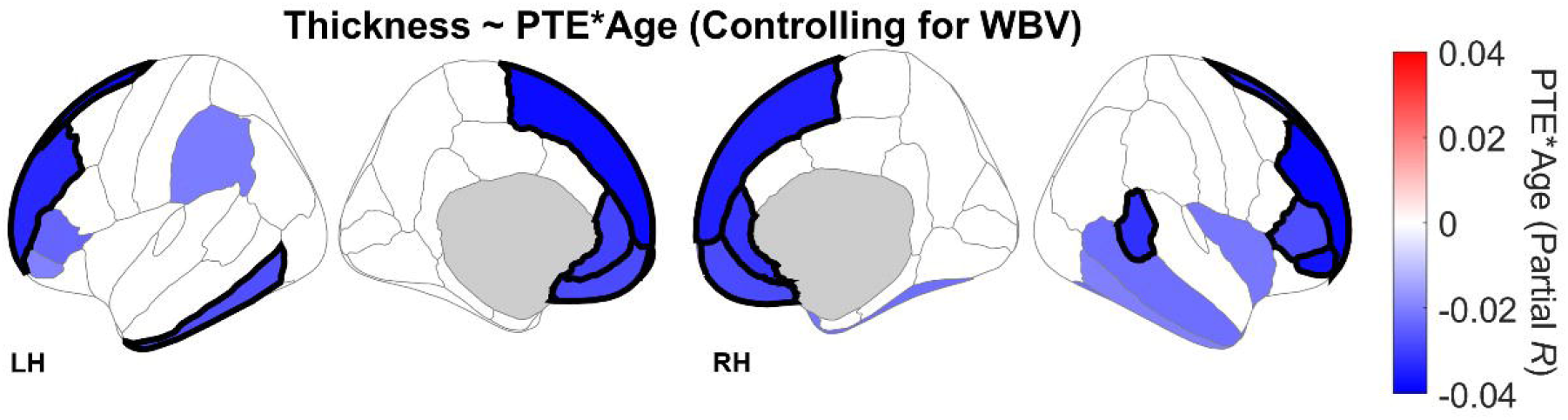
Prenatal tobacco exposure (PTE) × Age interactions on cortical thickness. Regions are color coded with respect to the color bar. Shaded regions indicate interactions that had *p*-values less than .05; regions that have thick borders indicate that those interactions also passed FDR correction. The degree of shading indicates greater negative (or lesser positive) associations with age in individuals with PTE. These images were generated in MATLAB using data from the *ggseg* toolbox in R.^42^ LH = left hemisphere. RH = right hemisphere. WBV = whole-brain volume.

#### PAE×PTE interactions

There were no FDR-corrected PAE×PTE or PAE×PTE×Age interactions on thickness (eTables 20-21) or surface area (eTables 27-28).

### Post Hoc Analyses

Post hoc analyses probed (1) the effects/interactions of PTE on cortical thickness/thinning after learning of the pregnancy and (2) the effects/interactions of PTE on cortical thickness/thinning when excluding individuals with PAE.

#### Knowing-of-Pregnancy PTE

These analyses included participants of caregivers reporting PTE after learning of the pregnancy (*n*=262) and the same 4,678 non-PTE participants from the initial analyses; here, between 10 and 214 participants were removed from individual-ROI analyses due to outlying/missing data. Model equations were identical to the initial analyses above. Collapsed across timepoints, PTE was associated with thinner bilateral-parahippocampal cortices, *p*s≤.038, and thicker bilateral-lingual cortices, *p*s≤.030, but these effects did not pass FDR correction (eTable 45). There were PTE×Age interactions in 5 regions (left-frontal-pole, right-inferior-parietal, right-pars-orbitalis, left-superior-frontal, left-rostral-anterior-cingulate), *p*s≤.039, none passing FDR correction (eTable 47). There were no FDR-corrected PAE×PTE or PAE×PTE×Age interactions (eTables 48-49).

#### PTE without PAE

These analyses included 3,917 participants whose caregivers did not report alcohol use during pregnancy (PTE: *n*=366, regardless of before/after learning of pregnancy; non-PTE: *n*=3,551); between 9 and 163 participants were removed from individual-ROI analyses due to outlying/missing data. Model equations were identical to the initial analyses except PAE-included factors were removed. Across timepoints, PTE was associated with thinner cortices in 4 regions (bilateral-parahippocampal, right-precentral, left-lateral-orbitofrontal), *p*s≤.048, with only the effect on right-parahippocampal thickness passing FDR correction; in 6 regions, PTE was associated with thicker cortices (bilateral-cuneus, right transverse-temporal, left-pars-opercularis, right-superior-temporal, right-temporal-pole), *p*s≤.039, which did not pass FDR correction (eTable 51). There were PTE×Age interactions in 9 regions (bilateral-inferior-temporal, bilateral-rostral-anterior-cingulate, bilateral-rostral-middle-frontal, left-frontal-pole, right-medial-orbitofrontal, left-superior-frontal), *p*s≤.036; only the association of faster cortical thinning of the left-frontal-pole in the PTE group passed FDR correction (eTable 52).

## Discussion

We described PAE and PTE moderation of early adolescent longitudinal cortical development. While there were associations with PAE, PTE’s longitudinal age effects were stronger and regionally broader. Notably, PTE was associated faster rates of frontal-lobe cortical thinning (Figure 3). Interestingly, in post hoc analyses (with their inherently reduced sample sizes given added exclusionary criteria), these effects were weaker, possibly due to systematic underreporting of PTE, differences in developmental timing, or loss of statistical power.

We know of two reports on PTE’s and/or PAE’s effects on cortical structure at multiple timepoints in ABCD.^12,18^ In one,^12^ participants were grouped based on responses to PTE/Before and PTE/After pregnancy recognition; cortical surface-area and volume data (collapsed across hemispheres) were analyzed separately at the baseline and two-year appointments. At both, they reported smaller precentral surface area in the PTE group, comparable to our results showing reduced bilateral-precentral area given PTE (albeit non-FDR-corrected) (Figure 2B, eTable 24). That report also described reduced area in entorhinal and postcentral cortices at baseline, while our results (which accounted for data at both timepoints) showed non-FDR-corrected reductions in the right-entorhinal and left-postcentral cortices (eTable 24). The overlap in these results (despite different sample sizes and analytical covariates) corroborates the robustness of these effects, with our analyses adding brain hemispheric spatial detail.

In the second report,^18^ the effects of PAE and PTE (among other prenatal exposures) on thickness, area, and volume were analyzed separately at the two appointments. As in that report, we showed FDR-corrected thinner bilateral-parahippocampal cortices in those with PTE (Figure 2A), with there being additional partial overlap across the two sets of results in PTE’s effects on thickness. While that study reported significantly thicker right-pars-triangularis cortices (which we also showed; eTable 17), our results revealed significant PTE×Age interactions there and its surrounding regions in both hemispheres (eTable 19). Thus, by incorporating a longitudinal trajectory, we described a broader pattern of faster frontal-lobe cortical thinning in those with PTE (Figure 3). As noted above, PTE is associated with reduced cortical thickness, surface area, and/or volume across regions,^3,8-15^ which was partially corroborated here (i.e., some cortical regions were thicker and more expansive in those with PTE). However, analyzing the developmental trajectories of thickness and surface area here informs how PTE and/or PAE may impact adolescent neuroanatomical development long after such prenatal insults to the developing brain.

PAE’s more minimal effects relative to PTE were contrary to our hypotheses. While one ABCD study similarly showed minimal PAE effects on cortical thickness, that report showed broader, FDR-corrected increases in cortical area given PAE.^18^ Another ABCD publication reported greater baseline regional cortical volumes and areas in those with PAE.^31^ Interestingly, that latter report indicated that its analyses controlled for ICV in volumetric analyses. While it is unclear if ICV was included in those surface-area analyses, it is worth noting that had our final analyses controlled for ICV instead of total surface area, our results would have suggested that the effects of PAE on cortical area were equally widespread and significant (eTable 30), suggesting that covariate selection (which, here, was informed by developmental trajectory) may equally impact our understanding of the effects of other predictors of interest (e.g., PAE, PTE).^25^

In contrast to PTE’s moderated and direct associations in initial analyses, there were weaker effects in post hoc analyses. Importantly, these latter analyses included dramatically smaller samples of participants in the PTE comparison group (initial analyses: *n*=739; knowing-of-pregnancy PTE analyses: *n*=262; no-PAE PTE: *n*=366), reducing analytical power. While these analyses could suggest that continuing tobacco use after learning of the pregnancy restores/protects against tobacco’s neurotoxic effects from its use prior to learning of the pregnancy, it is more likely that the tobacco use after learning of the pregnancy (and, presumably, before knowing) was underreported, considering ABCD’s retrospective collection of these data ∼10 years after such tobacco use would have occurred.

While underreporting (via self-report) of tobacco use/smoking during pregnancy can be prevalent when measured prospectively,^32-34^ retrospective reports may be generally accurate and reliable,^35^ potentially due to the reduced stigma in reporting such events when *not* pregnant,^36^ with there being good reliability between self-reports years apart.^37-39^ Interestingly, one study, which compared prospective identification of during-pregnancy smoking versus its retrospective recall 14.5 years later, found that 9% of women (prospectively identified as smokers) retrospectively indicated having never smoked during pregnancy (or having smoked but not after learning of the pregnancy).^35^ In another study that compared medical records (midwives’ records at first antenatal visits) with retrospective recall of during-pregnancy smoking ∼11 years later, the highest proportion of discordant reports were those whose medical records indicated they smoked cigarettes but retrospectively reported they were nonsmokers.^36^ Thus, some of the weaker associations in the post hoc knowing-of-pregnancy PTE (relative to the omnibus PTE) analyses may be because those who did use tobacco after knowing of pregnancy did not report doing so when asked retrospectively in ABCD. Alternatively, some of the interactions from the omnibus analyses may have gone undetected in the post hoc analyses because the faster cortical thinning (within the omnibus analyses) had already occurred in those with knowing-of-pregnancy PTE. In 6-to 8-year-olds, the strongest effects of PTE on cortical thickness were in those with PTE throughout the pregnancy, with no effects of PTE when the pregnant person’s tobacco use ceased upon learning of the pregnancy^9^; thus, the effects of before-knowing-of-pregnancy PTE may take longer to manifest, as potentially observed here. Overall, the post hoc analyses do not diminish those of the initial analyses but provide insight for future observational-study analyses (e.g., the HEALthy Brain and Child Development Study^40^).

We recognize that there are opportunities to build on the present results. Given ABCD’s design, we cannot biochemically verify PTE or PAE (or lack thereof). However, given ABCD’s collection of its participants’ deciduous teeth, there are potentialities to extend our analyses with respect to cotinine biomarkers of PTE,^41^ which would enhance the temporal resolution of ABCD’s PTE-related queries of tobacco use before/after learning of the pregnancy. Similarly, as ABCD neuroimaging began at ages 9-10, we cannot (in ABCD) evaluate PTE’s and PAE’s neuroanatomical effects before ages 9-10 and, not yet, how PAE/PTE impacts cortical development in later adolescence. But, as with potential cotinine measurements, ABCD provides a rich resource to better understand how the prenatal, postnatal, and early-childhood environments affect brain development. Given the current longitudinal investigation of cortical change in early adolescence, we have provided key insight into how the prenatal environment may influence how the brain continues to develop well into childhood and adolescence.

## Acknowledgments

Data used in the preparation of this article were obtained from the Adolescent Brain Cognitive Development^SM^ (ABCD) Study (https://abcdstudy.org), held in the NIMH Data Archive (NDA). This is a multisite, longitudinal study designed to recruit more than 10,000 children age 9-10 and follow them over 10 years into early adulthood. The ABCD Study® is supported by the National Institutes of Health and additional federal partners under award numbers U01DA041048, U01DA050989, U01DA051016, U01DA041022, U01DA051018, U01DA051037, U01DA050987, U01DA041174, U01DA041106, U01DA041117, U01DA041028, U01DA041134, U01DA050988, U01DA051039, U01DA041156, U01DA041025, U01DA041120, U01DA051038, U01DA041148, U01DA041093, U01DA041089, U24DA041123, U24DA041147. A full list of supporters is available at https://abcdstudy.org/federal-partners.html. A listing of participating sites and a complete listing of the study investigators can be found at https://abcdstudy.org/consortium_members/. ABCD consortium investigators designed and implemented the study and/or provided data but did not necessarily participate in the analysis or writing of this report. This manuscript reflects the views of the authors and may not reflect the opinions or views of the NIH or ABCD consortium investigators. The ABCD data repository grows and changes over time. The ABCD data used in this report came from 10.15154/1523041.

## Notes

**Conflict of Interest Statement:** The authors declare no competing financial interests.

### Competing Interest Statement

The authors have declared no competing interest.

## References

1. Ferschmann L, Bos MGN, Herting MM, Mills KL, Tamnes CK. Contextualizing adolescent structural brain development: Environmental determinants and mental health outcomes. Current Opinion in Psychology. 2022;44:170–176. doi:10.1016/j.copsyc.2021.09.014

2. Marshall AT, Betts S, Kan EC, McConnell R, Lanphear BP, Sowell ER. Association of lead-exposure risk and family income with childhood brain outcomes. Nature Medicine. 2020;26:91–97. doi:10.1038/s41591-019-0713-y

3. Rivkin MJ, Davis PE, Lemaster JL, et al. Volumetric MRI study of brain in children with intrauterine exposure to cocaine, alcohol, tobacco, and marijuana. Pediatrics. 2008;121(4):741–750. doi:10.1542/peds.2007-1399

4. Thomas G, Gonneau G, Poole N, Cook J. The effectiveness of alcohol warning labels in the prevention of fetal alcohol spectrum disorder: A brief review International Journal of Alcohol and Drug Research. 2014;3(1):91–103. doi:10.7895/ijadr.v3i1.126

5. McCormack C, Hutchinson D, Burns L, et al. Prenatal alcohol consumption between conception and recognition of pregnancy Alcoholism: Clinical and Experimental Research. 2017;41(2):369–378. doi:10.1111/acer.13305

6. Qato DM, Zhang C, Gandhi AB, Simoni-Wastila L, Coleman-Cowger VH. Co-use of alcohol, tobacco, and licit and illicit controlled substances among pregnant and non-pregnant women in the United States: Findings from 2006 to 2014 National Survey on Drug Use and Health (NSDUH) data. Drug and Alcohol Dependence. 2020;206:107729. doi:10.1016/j.drugalcdep.2019.107729

7. Donald KA, Eastman E, Howells FM, et al. Neuroimaging effects of prenatal alcohol exposure on the developing human brain: A magnetic resonance imaging review. Acta Neuropsychiatrica. 2015;27(5):251–269. doi:10.1017/neu.2015.12

8. Bublitz MH, Stroud LR. Maternal smoking during pregnancy and offspring brain structure and function: Review and agenda for future research. Nicotine & Tobacco Research. 2012;14(4):388–397. doi:10.1093/ntr/ntr191

9. El Marroun H, Schmidt MN, Franken IHA, et al. Prenatal tobacco exposure and brain morphology: A prospective study in young children. Neuropsychopharmacology. 2014;39:792–800. doi:10.1038/npp.2013.273

10. Gonzalez MR, Uban KA, Tapert SF, Sowell ER. Prenatal tobacco exposure associations with physical health and neurodevelopment in the ABCD cohort. Health Psychology. 2023;42(12):856–867. doi:10.1037/hea0001265

11. Marshall AT, Bodison SC, Uban KA, et al. The impact of prenatal alcohol and/or tobacco exposure on brain structure in a large sample of children from a South African birth cohort. Alcoholism: Clinical and Experimental Research. 2022;doi:10.1111/acer.14945

12. Puga TB, Dai HD, Wang Y, Theye E. Maternal tobacco use during pregnancy and child neurocognitive development. JAMA Network Open. 2024;7(2):e2355952. doi:10.1001/jamanetworkopen.2023.55952

13. Rivera PJR, Liang H, Isaiah A, et al. Prenatal tobacco exposure on brain morphometry partially mediated poor cognitive performance in preadolescent children. NeuroImmune Pharmacology and Therapeutics. 2023;2(4):375–386. doi:10.1515/nipt-2023-0013

14. Toro R, Leonard G, Lerner JV, et al. Prenatal exposure to maternal cigarette smoking and the adolescent cerebral cortex. Neuropsychopharmacology. 2008;33:1019–1027. doi:10.1038/sj.npp.1301484

15. Zou R, Boer OD, Felix JF, et al. Association of maternal tobacco use during pregnancy with preadolescent brain morphology among offspring. JAMA Network Open. 2022;5(8):e2224701. doi:10.1001/jamanetworkopen.2022.24701

16. Froggatt S, Covey J, Reissland N. Infant neurobehavioural consequences of prenatal cigarette exposure: A systematic review and meta-analysis. Acta Paediatrica. 2020;109(6):1112–1124. doi:10.1111/apa.15132

17. Jernigan TL, Brown SA, Dowling GJ. The Adolescent Brain Cognitive Development Study. Journal of Research on Adolescence. 2018;28(1):154–156. doi:10.1111/jora.12374

18. Gu Z, Barch DM, Luo Q. Prenatal substance exposure and child health: Understanding the role of environmental factors, genetics, and brain development. PNAS Nexus. 2024;3(1):pgae003. doi:10.1093/pnasnexus/pgae003

19. Garavan H, Bartsch H, Conway K, et al. Recruiting the ABCD sample: Design considerations and procedures. Developmental Cognitive Neuroscience. 2018;32:16–22. doi:10.1016/j.dcn.2018.04.004

20. Hagler DJ, Hatton SN, Cornejo MD, et al. Image processing and analysis methods for the Adolescent Brain Cognitive Development Study. NeuroImage. 2019;202:116091. doi:10.1016/j.neuroimage.2019.116091

21. Fischl B, Dale AM. Measuring the thickness of the human cerebral cortex from magnetic resonance images. Proceedings of the National Academy of Sciences. 2000;97(20):11050–11055. doi:10.1073/pnas.200033797

22. Dale AM, Fischl B, Sereno MI. Cortical surface-based analysis. I. Segmentation and surface reconstruction. NeuroImage. 1999;9(2):179–194. doi:10.1006/nimg.1998.0395

23. Talati A, Wickramaratne PJ, Wesselhoeft R, Weissman MM. Prenatal tobacco exposure, birthweight, and offspring psychopathology. Psychiatry Research. 2017;252:346–352. doi:10.1016/j.psychres.2017.03.016

24. de Kieviet JF, Zoetebier L, van Elburg RM, Vermeulen RJ, Oosterlaan J. Brain development of very preterm and very low-birthweight children in childhood and adolescence: A meta-analysis. Developmental Medicine & Child Neurology. 2012;54(4):313–323. doi:10.1111/j.1469-8749.2011.04216.x

25. Marshall AT, Adise S, Kan EC, Sowell ER. Longitudinal sex-at-birth and age analyses of cortical structure in the ABCD Study®. bioRxiv. 2024;doi:10.1101/2024.06.10.598367

26. Mills KL, Siegmund KD, Tamnes CK, et al. Inter-individual variability in structural brain development from late childhood to young adulthood. NeuroImage. 2021;242:118450. doi:10.1016/j.neuroimage.2021.118450

27. Leys C, Ley C, Klein O, Bernard P, Licata L. Detecting outliers: Do not use standard deviation around the mean, use absolute deviation around the median. Journal of Experimental Social Psychology. 2013;49(4):764–766. doi:10.1016/j.jesp.2013.03.013

28. Nakagawa S, Cuthill IC. Effect size, confidence interval and statistical significance: A practical guide for biologists. Biological Reviews. 2007;82:591–605. doi:10.1111/j.1469-185X.2007.00027.x

29. Aloe AM, Thompson CG. The synthesis of partial effect sizes. journal of the Society for Social Work and Research. 2013;4(4):390–405. doi:10.5243/jsswr.2013.24

30. Benjamini Y, Hochberg Y. Controlling the false discovery rate: A practical and powerful approach to multiple testing. Journal of the Royal Statistical Society Series B (Methodological). 1995;57(1):289–300.

31. Lees B, Mewton L, Jacobus J, et al. Association of prenatal alcohol exposure with psychological, behavioral, and neurodevelopmental outcomes in children from the Adolescent Brain Cognitive Development Study. The American Journal of Psychiatry. 2020;177(11):1060–1072. doi:10.1176/appi.ajp.2020.20010086

32. Boyd NR, Windsor RA, Perkins LL, Lowe JB. Quality of measurement of smoking status by self-report and saliva cotinine among pregnant women. Maternal and Child Health Journal. 1998;2:77–83. doi:10.1023/A:1022936705438

33. Shipton D, Tappin DM, Vadiveloo T, Crossley A, Aitken DA, Chalmers J. Reliability of self reported smoking status by pregnant women for estimating smoking prevalence: A retrospective, cross sectional study. BMJ. 2009;339:b4347. doi:10.1136/bmj.b4347

34. Owen L, McNeill A. Saliva cotinine as indicator of cigarette smoking in pregnant women. Addiction. 2002;96(7):1001–1006. doi:10.1046/j.1360-0443.2001.96710019.x

35. Pickett KE, Kasza K, Biesecker G, Wright RJ, Wakschlag LS. Women who remember, women who do not: A methodological study of maternal recall of smoking in pregnancy. Nicotine & Tobacco Research. 2009;11(10):1166–1174. doi:10.1093/ntr/ntp117

36. Post A, Gilljam H, Bremberg S, Galanti MR. Maternal smoking during pregnancy: A comparison between concurrent and retrospective self-reports. Paediatric and Perinatal Epidemiology. 2008;22:155–161. doi:10.1111/j.1365-3016.2007.00917.x

37. Jaspers M, de Meer G, Verhulst FC, Ormel J, Reijneveld SA. Limited validity of parental recall on pregnancy, birth, and early childhood at child age 10 years. J Clin Epidemiol. 2010;63(2):185–191. doi:10.1016/j.jclinepi.2009.05.003

38. Brigham J, Lessov-Schlaggar CN, Javitz HS, et al. Validity of recall of tobacco use in two prospective cohorts. American Journal of Epidemiology. 2010;172(7):828–835. doi:10.1093/aje/kwq179

39. Jacobson SW, Chiodo LM, Sokol RJ, Jacobson JL. Validity of maternal report of prenatal alcohol, cocaine, and smoking in relation to neurobehavioral outcome. Pediatrics. 2002;109(5):815–825. doi:10.1542/peds.109.5.815

40. Volkow ND, Gordon JA, Bianchi DW, et al. The HEALthy Brain and Child Development Study (HBCD): NIH collaboration to understand the impacts of prenatal and early life experiences on brain development. Developmental Cognitive Neuroscience. 2024;69:101423. doi:10.1016/j.dcn.2024.101423

41. Dolatmoradi M, Ellis J, Austin C, Arora M, Vertes A. Detection and imaging of exposure-related metabolites and xenobiotics in hard tissues by laser sampling and mass spectrometry. Analytical Chemistry. 2024;96:7022–7029. doi:10.1021/acs.analchem.4c00224

42. Mowinckel AM, Vidal-Piñeiro D. Visualisation of Brain Statistics with R-packages ggseg and ggseg3d. arXiv preprint arXiv:191208200. 2019;

